# Tobramycin adaptation enhances policing of social cheaters in *Pseudomonas aeruginosa*

**DOI:** 10.1101/2020.11.15.383836

**Authors:** Rhea G. Abisado, John H. Kimbrough, Brielle M. Mckee, Vaughn D. Craddock, Nicole E. Smalley, Ajai A. Dandekar, Josephine R. Chandler

## Abstract

The *Pseudomonas aeruginosa* LasR-I quorum sensing (QS) system regulates secreted proteases that can be exploited by cheaters, such as QS receptor-defective (*lasR*) mutants. *lasR* mutants emerge in populations growing on casein as a sole source of carbon and energy and increase in the population because they do not incur the substantial cost of engaging in QS. QS also increases resistance to some antibiotics, such as tobramycin. Here, we show that tobramycin suppresses the emergence of *lasR* mutants in casein-passaged populations. We also identify several mutations that accumulate in those populations indicating evidence of antibiotic adaptation. Mutations in one gene, *ptsP*, increase activity of the LasR-I system and production of a QS-controlled phenazine, pyocyanin. We find that mutations in *ptsP* lead to suppression of cheaters independent of tobramycin. Cheater suppression relies on pyocyanin, which acts as a policing toxin by targeting cheaters. These results show that tobramycin suppresses *lasR* mutants through two mechanisms: first, by directly acting on tobramycin-susceptible cheaters and second, by selecting mutations in *ptsP* that lead to pyocyanin-dependent policing. This work demonstrates how adaptive mutations can alter the dynamics of cooperator-cheater relationships, which might be important for populations adapting to antibiotics during infections.

## INTRODUCTION

Many Proteobacteria have quorum-sensing (QS) systems that sense and respond to *N*-acyl-homoserine lactones (AHLs) to cause population density-dependent changes in gene expression (1–3). AHL systems involve a LuxR-family signal receptor and a LuxI-family signal synthase (1, 4, 5). LuxR-I systems in many Proteobacteria control production of exoproducts such as proteases and toxins, which can be considered public goods (3, 6, 7). Public goods can be used by any member of the population; however, some population members do not contribute to public good production and are called freeloaders or cheaters (7–9). An example of a cheater is individuals with null mutations in the gene coding for the LuxR-family signal receptor; these mutants do not produce QS-dependent public goods but can still exploit public goods produced by QS cooperators in the population (8–10).

Cheater proliferation presents a serious threat to the stability of cooperating populations. Because public goods are metabolically costly, cheaters can overrun cooperators (8, 9, 11). In conditions where the public goods are required for growth, uncontrolled cheating can lead to population collapse (7, 12–14). Mechanisms must exist to control cheating in order for cooperative phenotypes to be maintained. One such mechanism is policing (15, 16). In bacteria, cooperators police cheaters by linking toxin and toxin-resistance factor production, such as through co-regulation of each of these by QS (16–18). QS can also control cheating through co-regulation of public goods with private goods (13) that benefit only producing cells. Previously, we showed that the soil bacterium *Chromobacterium subtsugae* (formerly *Chromobacterium violaceum*) uses QS to co-regulate a publicly available secreted protease with a privately available cell-associated tetracycline-specific efflux pump (10). When *C. subtsugae* is grown on casein as a sole carbon and energy source, QS-dependent protease production can be exploited by QS-defective cheaters. However, these cheaters are suppressed when tetracycline is included in the casein medium, because they do not express the efflux pump conferring resistance (10). This type of cheater suppression likely requires some other selective force ensuring that public and private goods are maintained under co-regulation (19).

In this study, we sought to determine if antibiotics can restrain the emergence of QS-defective cheaters in bacteria other than *C. subtsugae*. We were also interested in understanding how adaptation under antibiotic selection can alter the dynamics of cooperation and cheating. Previous reports show that AHL QS regulates antibiotic resistance in *Pseudomonas aeruginosa* (20–23). There are two AHL QS systems in *P. aeruginosa*, the LasR-I and RhlR-I systems. These two systems produce and respond to the signals 3-oxododecanoyl-homoserine lactone (3OC12-HSL) and butanoyl-homoserine lactone (C4-HSL), respectively (3, 24, 25). The systems are hierarchical with LasR-I controlling activation of RhlR-I (25, 26). In biofilm conditions, deletions in LasR were previously shown to cause sensitivity to tobramycin antibiotic in at least one strain of *P. aeruginosa*, PAO1(20–22).

Here, we show the LasR-I system increases tobramycin resistance in planktonic conditions in the *P. aeruginosa* strain PA14. We also show that tobramycin can suppress the emergence of cheaters in cooperating PA14 populations grown on casein, similar to previous observations with *C. subtsugae* (10). We sequenced the genomes of isolates from tobramycin-evolved populations. All of the isolates had mutations in or upstream of *ptsP*, a gene coding for phosphoenolpyruvate-protein phosphotransferase (EI^Ntr^). This enzyme is the first in a global regulatory system known as the nitrogen phosphotransferase system (PTS^Ntr^) (27). Mutations in *ptsP* were previously shown to increase LasR-I activity through a mechanism that is not well understood (28). Interestingly, we observed that *ptsP* mutations suppress cheaters, even in populations passaged with no antibiotic. We demonstrate the cheater suppression is due to the QS-controlled toxin pyocyanin, which polices cheaters by selectively blocking their proliferation. Our results show that genetic adaptation to tobramycin can enhance policing against social cheaters. These results provide new information on policing mechanisms in *P. aeruginosa* and demonstrate how antibiotic selection can lead to changes in cooperative activity.

## MATERIALS AND METHODS

### Culture conditions and reagents

Bacteria were routinely grown in Luria-Bertani broth (LB) or LB buffered to pH 7 with 50 mM 3-(morpholino)-propanesulfonic acid (MOPS), or on LB agar (LBA; 1.5% (weight per volume) Bacto-Agar). Growth media for specific experiments were M9-caseinate (casein broth; 6 g l^-1^ Na_2_HPO_4_, 3 g l^-1^ KH_2_PO_4_, 0.5 g l^-1^ NaCl, 1 g l^-1^ NH_4_Cl; pH 7.4, 1% sodium caseinate); MOPS minimal medium (25 mM D-glucose, freshly prepared 5 μM FeSO_4_, 15 mM NH_4_Cl, and 2 mM K_2_HPO_4_ added to a 1X MOPS base buffer consisting of 50 mM MOPS, 4 mM tricine, 50 mM NaCl, 1 mM K2SO4, 50 μM MgCl2, 10 μM CaCl_2_, 0.3 μM (NH_4_)_6_Mo_7_O_24_, 40 μM H_3_BO_3_, 3 μM Co(OAc)_2_, 1 μM CuSO_4_, 8 μM MnSO_4_, and 1 μM ZnSO_4_) (29); a modified M9-casamino acid broth (200 ml l^-1^ 10X M9, 0.6 g l^-1^ thiamine hydrochloride, 1 ml l^-1^ 1 M MgSO_4_, 1 ml l^-1^ 0.2 M CaCl_2_, 10 g l^-1^ casamino acid; freshly prepared; filter sterilized), Pyocyanin Producing Media (PPM) (30), and 4% skim milk agar (SMA) (9). All *P. aeruginosa* broth cultures were grown at 37°C with shaking at 250 rpm, 18 mm test tubes (for 2 ml cultures), 125 ml baffled flasks (10 ml cultures), or 250 ml baffled flasks (60 ml cultures), unless otherwise specified. For *E. coli*, 100 μg ml^-1^ carbenicillin, 15 μg ml^-1^ gentamicin, and 10 μg ml^-1^ tetracycline were used. For *P. aeruginosa*, 300 μg ml^-1^ carbenicillin, 50–200 μg ml^-1^ gentamicin, and 200 μg ml^-1^ tetracycline were used. AHLs were purchased from Cayman Chemicals (MI, USA) and handled as described elsewhere (10). Genomic or plasmid DNA was extracted using Qiagen Puregene Core A kit (Hilden, Germany) or IBI Scientific plasmid purification mini-prep kit (IA, USA) while PCR products were purified using IBI Scientific PCR clean-up/gel extraction kits, according to the manufacturer’s protocol. All antibiotics were purchased from GoldBio (MO, USA) except for tetracycline which is from Fisher Scientific (PA, USA). Pyocyanin and DMSO (solvent for pyocyanin) were purchased from Sigma Aldrich (MO, USA) and Acros Organics (WI, USA), respectively.

### Bacterial strains and strain construction

All bacterial strains, plasmids, and primers used in this study are listed in Tables S1–S3. *P*. *aeruginosa* strain UCBPP-PA14 (‘PA14’) and PA14 derivatives were used for these studies. Markerless deletions in specific loci of *P*. *aeruginosa* PA14 were generated using allelic exchange as described previously (31). To generate plasmids for allelic exchange, DNA fragments with the mutated or deleted gene allele plus 500 bp flanking DNA were generated by PCR using primer-incorporated restriction enzyme sites. The PCR product was digested and ligated to pEXG2 *fusA1* G1634A) or pEX18Ap (Δ*lasR* and Δ*rhlR*) and transformed into the appropriate *P. aeruginosa* strain. The plasmids for Δ*lasI* and Δ*ptsP* were described elsewhere (27, 32). Merodiploids were selected on Pseudomonas Isolation Agar (PIA)-carbenicillin (300 μg ml^-1^) for Δ*lasR* and Δ*rhlR*, PIA-gentamicin (200 μg ml^-1^) for Δ*lasI* and Δ*ptsP*, and LBA-gentamicin (50 μg ml^-1^) for *fusA1* G1634A. Deletion mutants were counterselected using NaCl free-15% sucrose. Putative mutants were verified through antibiotic sensitivity tests and gene-targeted Sanger sequencing. Plasmid transformations were described previously (27, 33–35). Complementation strains were constructed by integrating the mini□CTX□1 vector at the neutral chromosomal *attB* locus (27, 34).

### Evolution experiments

To prepare the inoculum for evolution experiments, overnight (18 h) pure cultures were grown in LB-MOPS, diluted 1:50 into 2 ml LB-MOPS, and grown to an OD600 of ~3.5. To start the experiment, 50 μl from this starter culture was transferred to 2 ml fresh casein broth in an 18 mm tube. At 24 h intervals, cultures were diluted 1:50 into fresh casein broth in a new tube. Tobramycin was added every other day where indicated, similar to previous experiments with *C. subtsugae* (10). CFU ml^-1^ for each lineage was determined by viable plate counts every 96 h. The % *lasR* mutant cheater (*lasR* cheater) was determined by patching 100 colonies, unless otherwise specified, on SMA. Sixteen *lasR* mutants were sequence-verified by Sanger sequencing (Table S4).

### Antimicrobial susceptibility assay

Tobramycin susceptibility was determined by MIC according to the 2020 guidelines of the Clinical and Laboratory Standards Institute (CSLI), using a modified dilution method. Briefly, tobramycin was added to MOPS minimal medium and successively diluted 2-fold in a 200 μl volume in 2 ml tubes. For each experiment, two dilution series were staggered by starting them at different tobramycin concentrations to cover a broader range of concentrations. *P. aeruginosa* inocula were from LB-grown stationary-phase cells (OD_600_ of 3.2–4), being careful to ensure that all cultures in each experiment were grown to an identical cell density. The cultures were diluted 1:40 into each tube containing tobramycin, and the tubes were incubated with shaking for 20 h. After incubation, turbidity was measured using a BioTek Synergy 2 plate reader. The MIC was defined as the lowest concentration of tobramycin (μg ml^-1^) in which bacterial growth was not measurable. In some cases, CFU ml^-1^ was also determined by viable plate counts.

To determine susceptibility to *P. aeruginosa* culture fluid, we prepared culture fluid by inoculating overnight cultures to an optical density at 600 nm (OD_600_) of 0.1 into 60 ml casein broth, grew these cultures for 20 h, then passed cultures through a 0.22 μm filter to remove cells. The filtered fluid was mixed with 100 μl M9-casamino acid broth to a final volume of 2 ml and this was inoculated at an initial OD_600_ of 0.001 with either wild type, Δ*lasR, ΔptsP*, or Δ*ptsP* Δ*lasR* from logarithmic-stage LB-MOPS cultures at an OD600 0.2–0.6. The M9-casamino-filtrate mixture was incubated for 24 h and the initial and final population counts (CFU ml^-1^) were enumerated by colony counting on plates.

### Whole genome sequencing

Genomic DNA was extracted using the Qiagen Puregene yeast/bacteria kit and a sequencing library was constructed with 350-bp inserts (strain T2) or 200-bp inserts (all other strains). Sequencing was performed using Illumina HiSeq 4000 (for strain T2) or Illumina MiSeq with ~25X coverage (all other strains). The raw reads were aligned to the *P. aeruginosa* UCBPP-PA14 reference genome (UCBPP-PA14 Accession NC_008463) using Strand NGS (Bangalore, India) software v 3.1.1, using a pipeline described previously (36). Mutations of interest were verified by gene-targeted Sanger sequencing. Sequence reads for the ancestral PA14 (SAMN16823471) and tobramycin-evolved isolates (SAMN16823472 - SAMN16823478) can be found at the NCBI SRA under BioProject PRJNA678537.

#### Measurements of LasR activity and pyocyanin production

To measure LasR activity, we first introduced the LasR-responsive plasmid pBS351 to *P. aeruginosa* mutants or wild-type PA14 strains by electroporation (35). Electrocompetent cells were prepared from overnight cultures using 300 mM sucrose (37). Transformants were selected on LB agar using gentamicin at 50–200 μg ml^-1^ and routinely grown with gentamicin (50 μg ml^-1^ for agar and 15 μg ml^-1^ for broth) for plasmid maintenance. Transformants were grown in LB-MOPS with 15 μg ml^-1^ gentamicin for 18 h, washed with phosphate buffered saline (PBS), and fluorescence was measured using a BioTek Synergy 2 plate reader. For pyocyanin measurements, we extracted pyocyanin as described previously (38–40). Briefly, cells were grown for 18 h in Pyocyanin-producing media (30), and 5 ml whole culture was extracted with 2 ml chloroform. The organic layer was separated and extracted a second time with 0.2 N HCl. The absorbance of the aqueous layer was measured at 520 nm and multiplied by 17.072 to calculate the pyocyanin concentration (μg ml^-1^) (39). LasR activity and pyocyanin measurements were normalized to culture density (optical density at 600 nm) for reporting data.

### Coculture assays

Coculture experiments were conducted in 2 ml casein broth in 18 mm test tubes. To prepare the inoculum, overnight (18 h) pure cultures were grown in LB-MOPS, diluted to an optical density at 600 nm (OD_600_) of 0.025 for cheaters or 0.05 for cooperators into LB-MOPS, and grown to an OD_600_ of ~3.5 before combining at a 99:1 (cooperator: cheater) ratio. This mixture was then diluted 1:40 into casein broth to start the coculture. Cocultures were diluted 1:40 into fresh casein broth in a new tube every 24 h until the end of the experiment at 72 h. The initial and final total population counts (CFU ml^-1^) were determined by viable plate counts. The % *lasR* cheater was determined by patching 200 colonies on SMA.

## RESULTS

### LasR promotes tobramycin resistance in *P. aeruginosa* PA14 planktonic cultures

As an initial test that LasR contributes to tobramycin resistance in planktonic conditions, we determined the minimum inhibitory concentration (MIC) of tobramycin against the laboratory strain PA14 or PA14 with a deletion of *lasR* or *lasI*. To increase the MIC detection sensitivity, we generated two tobramycin dilution series that were staggered by using two different starting concentrations (41). We observed a small but reproducible 1.7-fold decrease in the Δ*lasR* mutant MIC relative to PA14 (Fig. 1A). The difference was observed when tobramycin was used to treat cultures grown to an optical density at 600 nm of 4 (OD_600_ 4), but not cultures at lower cell densities (Fig. S1). After 24 h treatment with 1.1 μg ml^-1^ tobramycin, about 200-fold fewer Δ*lasR* mutant cells were recovered than wild type (Fig. 1B). We also observed a decrease in the MIC of the Δ*lasI* mutant similar to that of the Δ*lasR* mutant (Fig. 1A). We could restore resistance to the Δ*lasI* mutant by adding synthetic 3OC12-HSL (Fig. 1A). There was no significant difference between the wild type and the Δ*rhlR* mutant, supporting the hypothesis that the RhlR system is not important for the resistance phenotype. Further, the MIC of the Δ*lasR*, Δ*rhlR* double mutant was similar to that of the Δ*lasR* single mutant. Together, these results show that the LasR-I system, but not the RhlR-I system, contributes to tobramycin resistance in planktonically-grown *P. aeruginosa* strain PA14.

**Fig. 1.**
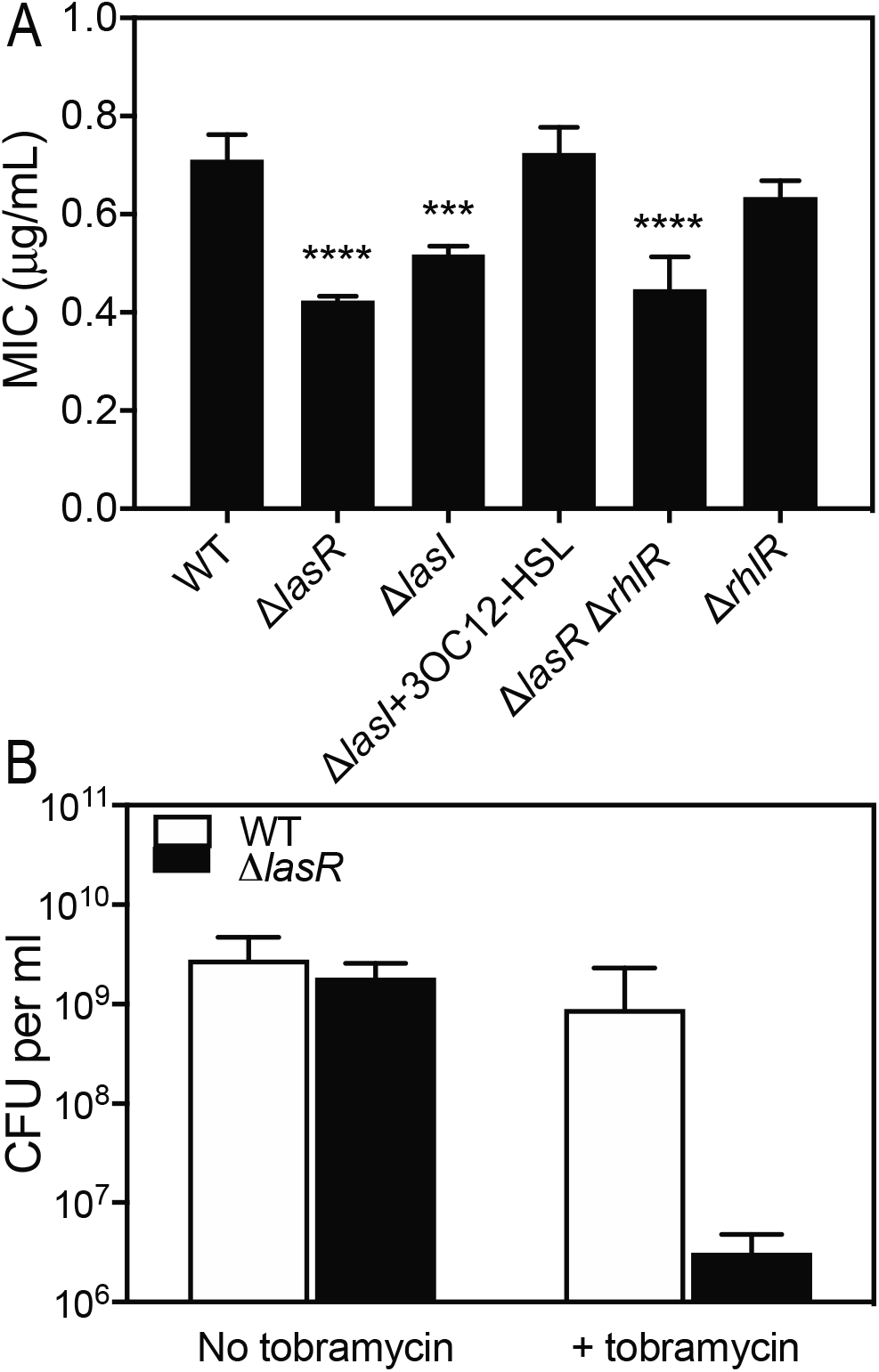
*lasR* contributes to tobramycin resistance in planktonically-grown *P. aeruginosa* PA14. **A**. The minimum inhibitory concentration (MIC) of tobramycin was determined for each of the strains indicated as described in Materials and Methods. 3OC12-HSL was added before inoculation where indicated (10 μM final). Statistical analysis by one-way ANOVA and Dunnett’s multiple comparisons test with wild type: *** p<0.001, **** p<0.0001. **B.** Cells recovered following tobramycin treatment (1.1 μg ml^-1^). Cells were treated as in part A and the surviving cells were enumerated by serial dilution and plating following treatment. For both experiments, the values shown represent the average of three independent experiments and the error bars represent the standard deviation.

### Tobramycin suppresses cheating in *P. aeruginosa*

*P. aeruginosa* requires a QS-controlled protease to grow in minimal medium with casein as the sole source of carbon and energy, which can be exploited by *lasR*-mutated cheaters that emerge in populations repeatedly passaged under these conditions (8, 9). We hypothesized that tobramycin might suppress *lasR* mutants in cooperating *P. aeruginosa* populations passaged on casein. To test this hypothesis, we assessed the frequency of *lasR* mutants in *P. aeruginosa* populations in casein broth or in broth with 0.6 μg ml^-1^ tobramycin, a concentration determined to be sub-MIC in these conditions. We distinguished *lasR* mutants by colony phenotypes (42, 43), which was confirmed in a subset of mutants by sequencing the *lasR* gene (Table S3). In populations with no antibiotic, *lasR* mutants emerged between 5–8 days and increased to >99% of the population (Fig. 2). None of our populations showed evidence of a collapse, as has been reported in similar experiments with PAO1 (9, 13, 16). In the populations with tobramycin, *lasR* mutant frequencies remained low or below the detection level throughout the experiment (Fig. 2). These results show that tobramycin can suppress the emergence of LasR-mutated cheaters in casein-grown *P. aeruginosa* populations.

**Fig. 2.**
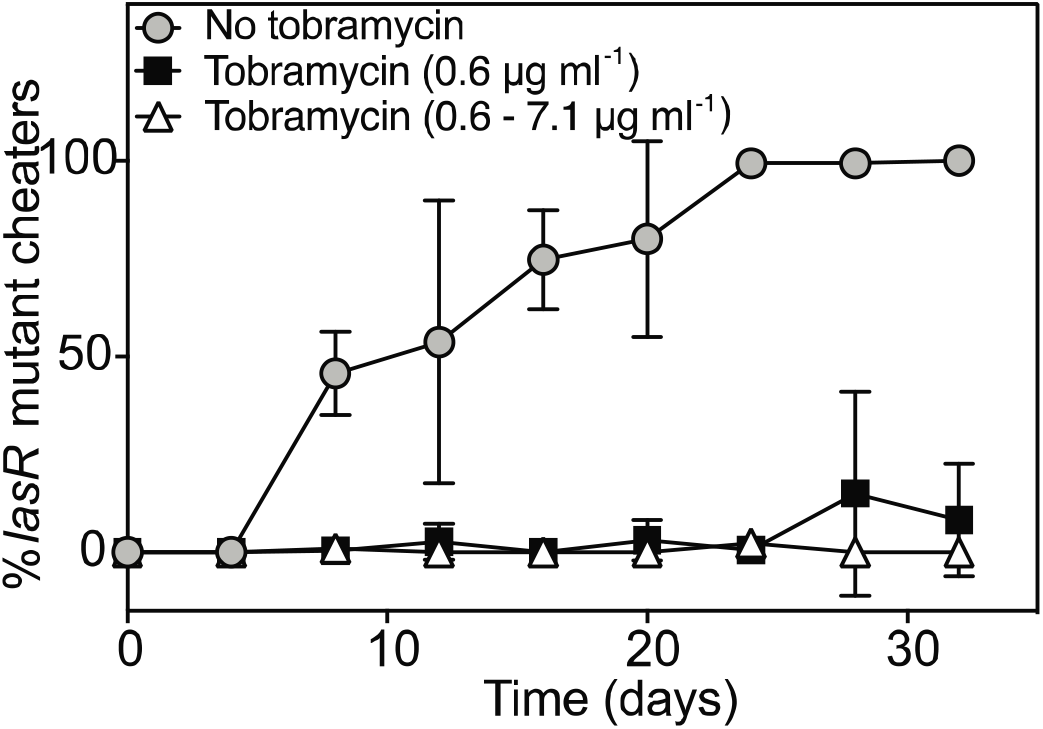
Tobramycin suppresses the emergence of *lasR*-mutant cheaters. *P. aeruginosa* populations were transferred daily in 1% casein broth for 32 days, and cheaters were enumerated every 4 days by patching as described in Materials and Methods. For each experiment, one culture was initially started with no tobramycin, and after 72 hr split into three cultures propagated in one of three conditions; i) with no tobramycin 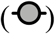, ii) with tobramycin added at 0.6 μg ml^-1^ 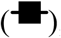, iii) with tobramycin added at 0.6 μg ml^-1^ initially, and increased by 50% every 4 days to a final concentration of 7.1 μg ml^-1^ 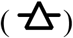. For conditions ii and iii, tobramycin was added every other day just after transfer to fresh medium. The detection level of cheaters was 1%. The values shown represent the average of three independent experiments and the error bars represent the standard deviation.

### Variants from tobramycin-treated populations undergo genetic adaptation

To assess how antibiotic adaptation influences cheater emergence and suppression, we carried out a second experiment using higher tobramycin concentrations for stronger selection. We used an initial concentration of 0.6 μg ml^-1^ tobramycin and increased the concentration by 50% every 4 days to reach a final concentration of 7.1 μg ml^-1^. In three independent cultures, the *lasR* mutant population remained at less than 5% of the total population (Fig. 2). We did not observe any significant growth inhibition at any stage of the experiment, even at the highest tobramycin concentration (Fig. S2), suggesting genetic adaptation occurred during passage. To test this hypothesis, we determined the MIC of one isolated variant from each of the passaged populations (variants T1, T2, and T3 from populations passaged with 0.6–7.1 μg ml^-1^ tobramycin and variants T4, T5, and T6 from populations passaged with 0.6 μg ml^-1^ tobramycin). All six variants showed a higher tobramycin MIC than the ancestor strain (Fig. S3A) or variants from identically treated populations with no added antibiotic (clones N1–N3, Fig. S3B).

To identify the mutations that accumulated in tobramycin-evolved variants, we sequenced the genomes of our six tobramycin-evolved variants. For comparison, we also sequenced the parent PA14 strain and clone N2 described above, an isolate from a population passaged with no antibiotic. We identified 3–6 mutations in each of the tobramycin-evolved variants that were not in either the parent PA14 or isolate N2 (Table 1 and S5). Most of the tobramycin-evolved variants had mutations in two genes: *ptsP*, which codes for phosphoenolpyruvate protein phosphotransferase and *fusA1*, which codes for translation elongation factor EF-G1A and is considered essential (44). To verify the role of *ptsP* and *fusA1* mutations in tobramycin resistance, we introduced mutations of each to the PA14 genome. We used Δ*ptsP* or the *fusA1* G1643A mutation from isolate T5, because *fusA1* deletions are thought to be nonviable (44). We also constructed a Δ*ptsP, fusA1* G1634A double mutant. We compared the MIC of the mutated strains with that of the PA14 parent (Fig. S3C and S3D). Each of the individual mutations increased PA14 resistance by about 2-fold and combining mutations increased the MIC by about 3-fold (Fig S3E).

**Table 1.**
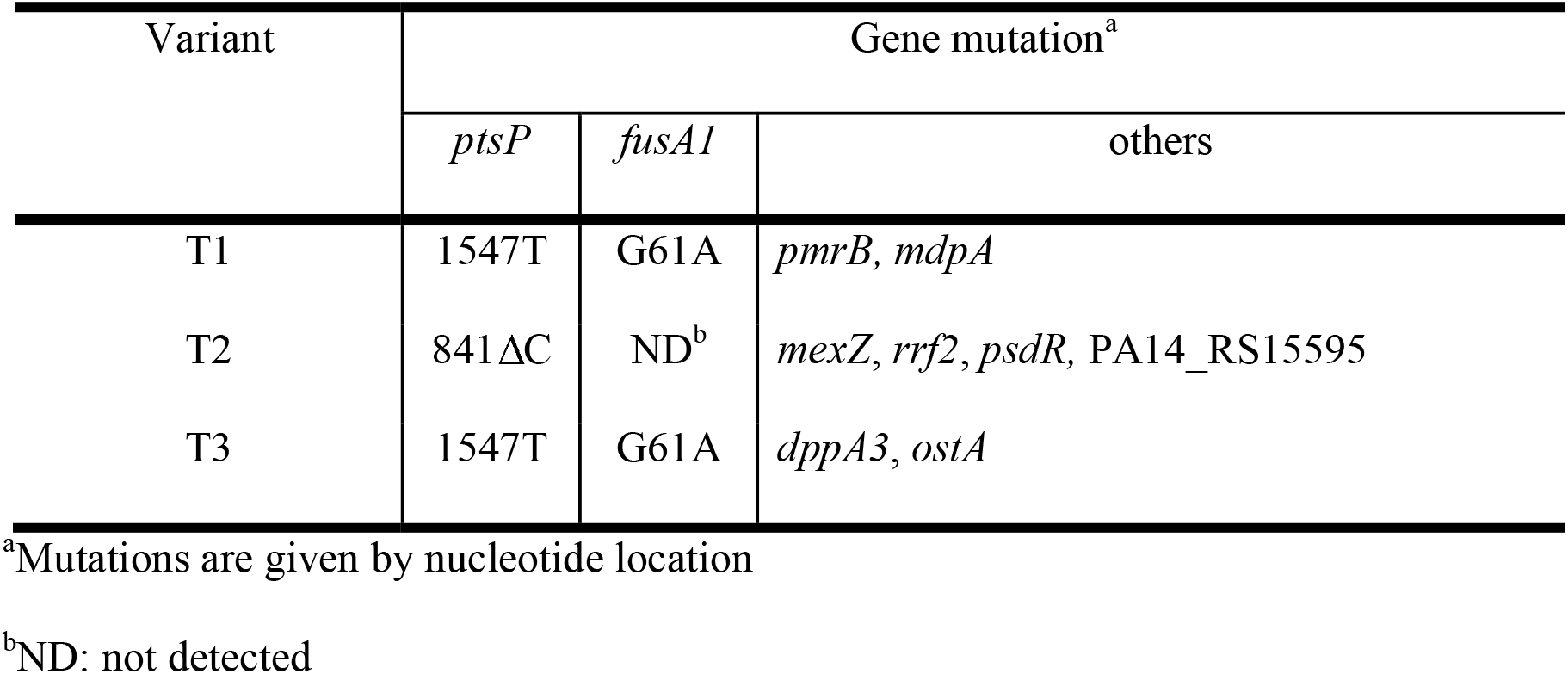
Mutations in variants from populations passed with 0.6-7.1 μg ml^-1^ tobramycin.

### *ptsP*-mutated strains have enhanced LasR activity and pyocyanin production

We focused our attention on the *ptsP* mutation because disrupting *ptsP* was previously shown to increase LasR activity (28), and we were interested in understanding how these effects could alter the cooperator-cheating dynamic. First, we tested the hypothesis that the LasR-LasI system is elevated at least in the T1, T2 and T3 mutants. To do so, we used a *PlasI-gfp* plasmid reporter (35), in which the *gfp* reporter gene is fused to the LasR-responsive *lasI* promoter. This reporter showed ~2–3-fold higher fluorescence in the Δ*ptsP* mutant than that of the wild-type PA14 strain (Fig. S4A), similar to previous results (28). The T1, T2 and T3 variants carrying this reporter also showed ~3-fold higher fluorescence than that of PA14. The elevated fluorescence levels in these variants could be restored to wild-type levels by introducing *ptsP* to the neutral *attB* site in the genome (Fig. 3A). The reporter activities of isolates N1, N2, and N3 were similar to that of PA14 (Fig. S4B). The difference in reporter activity in the PA14 Δ*ptsP* mutant or the T1–T3 strains could not be explained by differences in growth (Table S6).

**Fig. 3.**
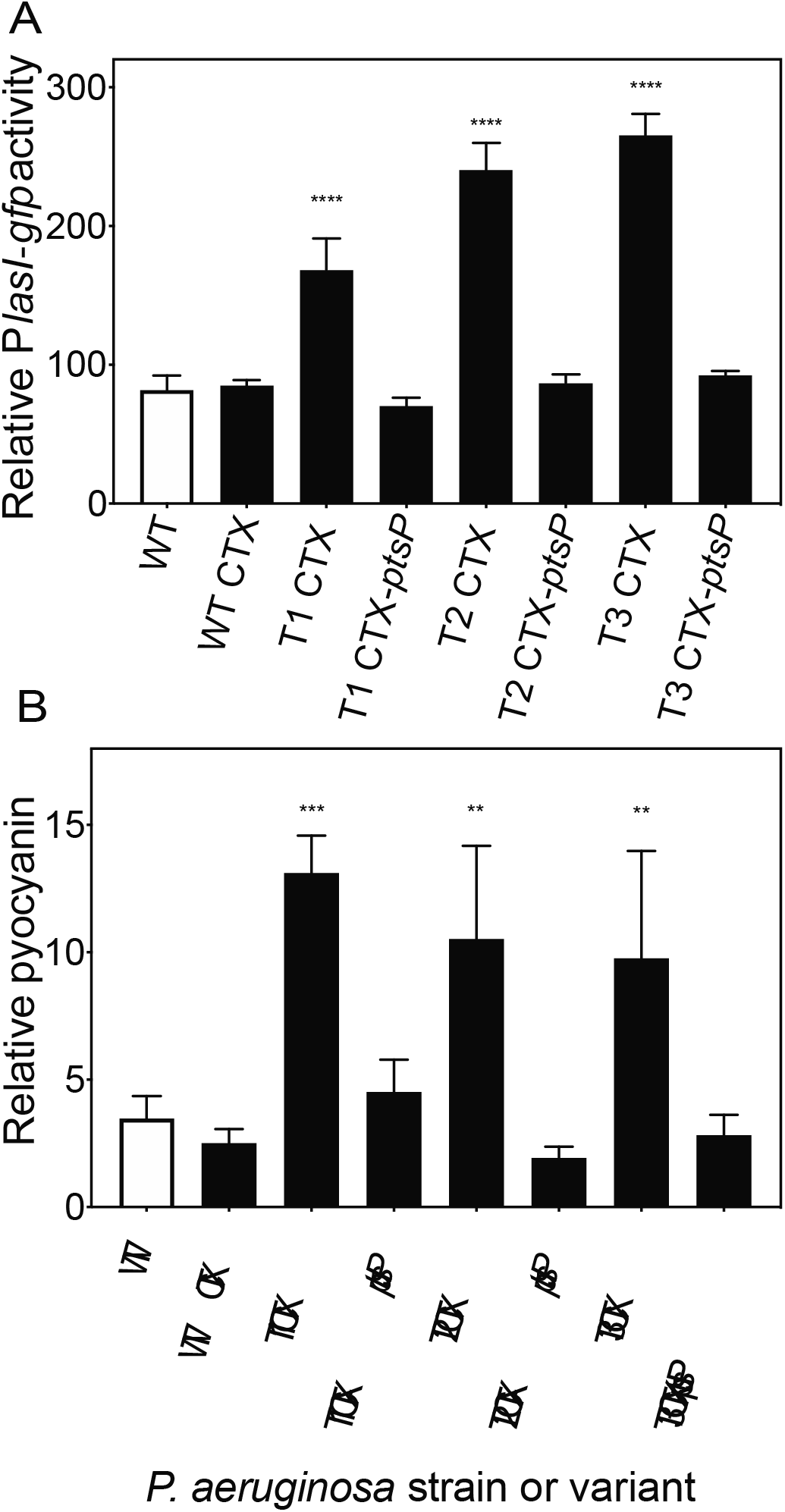
Effects of *ptsP* inactivation on LasR activity and pyocyanin production. **A.** Activity of LasR. *P. aeruginosa* strains were electroporated with a plasmid containing a LasR-responsive GFP reporter (pBS351). Reported values are fluorescence normalized to culture density at 18 h of growth. **B.** Pyocyanin production. Cultures were inoculated into pyocyanin-producing media, grown for 18 h, and extracted before quantifying pyocyanin as described in Materials and Methods. In all cases, reported values are μg/ml pyocyanin normalized to culture density at the time of measurement. Strains carried either the empty CTX-1 cassette (CTX), or the CTX-1-*ptsP* cassette, inserted at the neutral *attB* site in the genome. The values represent the average of three independent experiments and the error bars represent the standard deviation. Statistical analysis by one-way ANOVA and Dunnett’s multiple comparisons test with wild type: ** p<0.01, *** p<0.001, **** p<0.0001.

Disruptions in *ptsP* were previously shown to increase pyocyanin production by about 3-fold (28). To test whether pyocyanin production was similarly elevated in our tobramycin-evolved variants, we compared pyocyanin produced by the T1, T2, and T3 variants with that of PA14. Our results showed the variants have ~3-fold higher pyocyanin production than the PA14 parent, and that pyocyanin production can be reduced to wild-type levels in the variants by introducing *ptsP* to the *attB* site in the genome (Fig. 3B). This increase was not observed in isolates N1–N3 (Fig. S4C). Thus, *ptsP* mutations also increase pyocyanin production in strains passaged with tobramycin.

### *lasR*-mutant cheaters are suppressed in *ptsP* mutant populations

Because *ptsP* mutations alter QS regulation, we predicted that these mutations might influence cheating dynamics even in the absence of tobramycin. Specifically, we predicted *lasR* mutant cheaters might emerge at a faster rate in *ptsP* mutant populations due to the metabolic burden associated with the higher LasR activity. To test this hypothesis, we passaged the T1, T2, and T3 variants in casein broth for 32 days in the absence of antibiotic and monitored the emergence of *lasR*-mutant cheaters (Fig. 4A and S5). Surprisingly, cheaters emerged later in these populations compared with that of the PA14 ancestor (23 to >32 days for T1–T3 vs. 6–8 days for PA14). For T2, cheaters were never observed even by 32 days. Further, the total cheater frequency in the T1 and T3 populations did not reach the same levels as that of PA14 (33–67% for T1 and T3 vs. >98% for PA14) (Fig. S5A). There were no differences in cheating observed with variants N1–N3 passaged with no tobramycin, compared with PA14 (Fig. S5C). The total cell density remained similar for all evolution experiments, suggesting the differences in cheating were not due to differences in total population of the experiments (Fig. S5B, S5D).

**Fig. 4.**
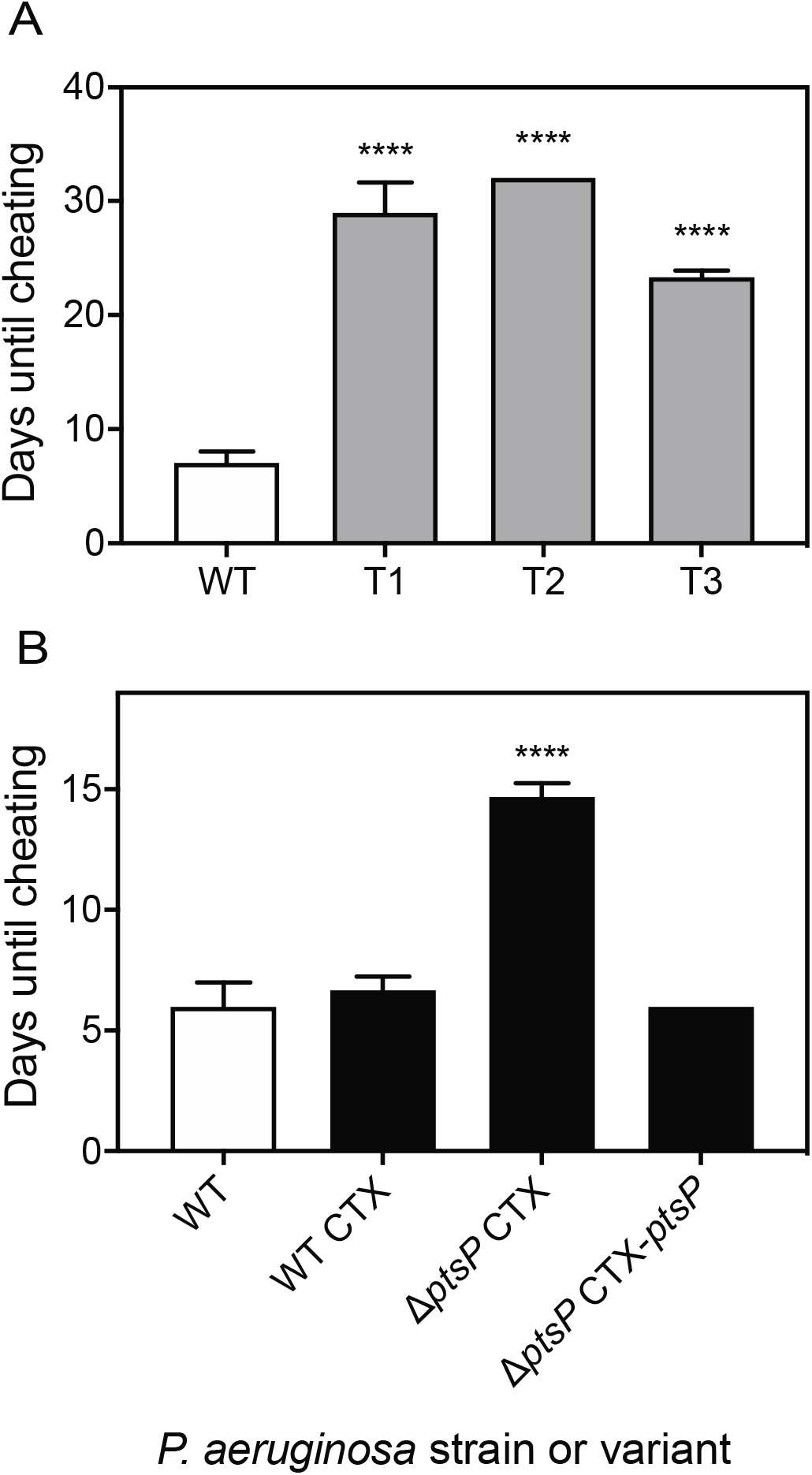
*ptsP* inactivation suppresses cheaters. P. *aeruginosa* populations were transferred daily in 1% casein broth for 32 days, and cheaters were enumerated every 4 days by patching as described in Materials and Methods. Values shown indicate the number of days until *1asR*-mutated cheaters emerge in **A**) populations of wild type PA14 (WT) and PA14 variants (T1, T2 and T3) from tobramycin-evolved populations and **B**) WT or the Δ*ptsP* mutant carrying either the empty CTX-1 cassette (CTX), or the CTX-1-*ptsP* (CTX-*ptsP*) cassette, inserted at the neutral *attB* site in the genome. Cheater emergence is defined as when *lasR* mutants reach a frequency in the population of 8%, at which point it was generally observed that cheater frequencies continued increasing rather than decreasing or remaining constant. Full data sets of cheater frequency and total populations are shown in Fig. S5. Values represent the average of three independent experiments and the error bars represent the standard deviation. Statistical analysis by one-way ANOVA and Dunnett’s multiple comparisons test with wild type for A; and by oneway ANOVA and Tukey’s multiple comparisons test for B: **** p<0.0001.

We hypothesized that the difference in cheating observed with the tobramycin-evolved variants was due to the disrupted *ptsP* gene in these strains. To test this hypothesis, we tested cheating in a Δ*ptsP* mutant. We passaged the Δ*ptsP* mutant carrying either CTX-1 or the CTX-1-*ptsP* cassette in 1% casein broth for 32 days and monitored cheater frequency over time. We also monitored cheating in PA14 and PA14 CTX-1. We observed no differences in cheating of PA14 CTX-1 compared with PA14, verifying that the CTX-1 cassette had no effect on cheating. However, cheaters emerged later and reached lower levels overall in the Δ*ptsP-CTX-1* populations compared with either PA14 strain (Fig. 4B, S5E). Cheating was indistinguishable from PA14 in the Δ*ptsP*-CTX-1-*ptsP* populations (Fig. 4B, S5E). There were no notable differences in total growth of any of these populations (Fig. S5F). Together, our results show that *ptsP* mutations cause delayed cheating and reduced cheater loads.

### Pyocyanin produced by Δ*ptsP* mutants is active against *lasR*-mutated cheaters

A potential explanation for delayed cheating in the Δ*ptsP* mutant populations is that the Δ*ptsP* mutant has a growth advantage compared with the PA14 ancestral strain. However, PA14 and Δ*ptsP* showed identical growth rates (Table S6), suggesting there is another explanation. We hypothesized that the Δ*ptsP* mutant produces a toxin that inhibits growth of *lasR* mutant cheaters. To test this hypothesis, we filtered culture fluid from a Δ*ptsP* mutant and tested its ability to inhibit growth of logarithmically growing PA14 or PA14 with deletions of *ptsP, lasR*, or both *ptsP* and *lasR*. We performed identical treatments with filtered fluid from PA14 or with unspent culture media (untreated control) and enumerated cell growth after 24 h. Treatment with PA14 culture fluid had no effects on growth and was similar to the untreated control (Fig. 5). However, treatment with the Δ*ptsP* mutant filtrate reduced growth of the Δ*lasR* mutant by 83-fold and Δ*ptsP* Δ*lasR* mutants by 13,000-fold, compared with no treatment. Thus, the Δ*ptsP* mutant secretes a substance that is growth inhibitive to *lasR*-mutated strains and particularly to Δ*ptsP* Δ*lasR* mutants.

**Fig. 5.**
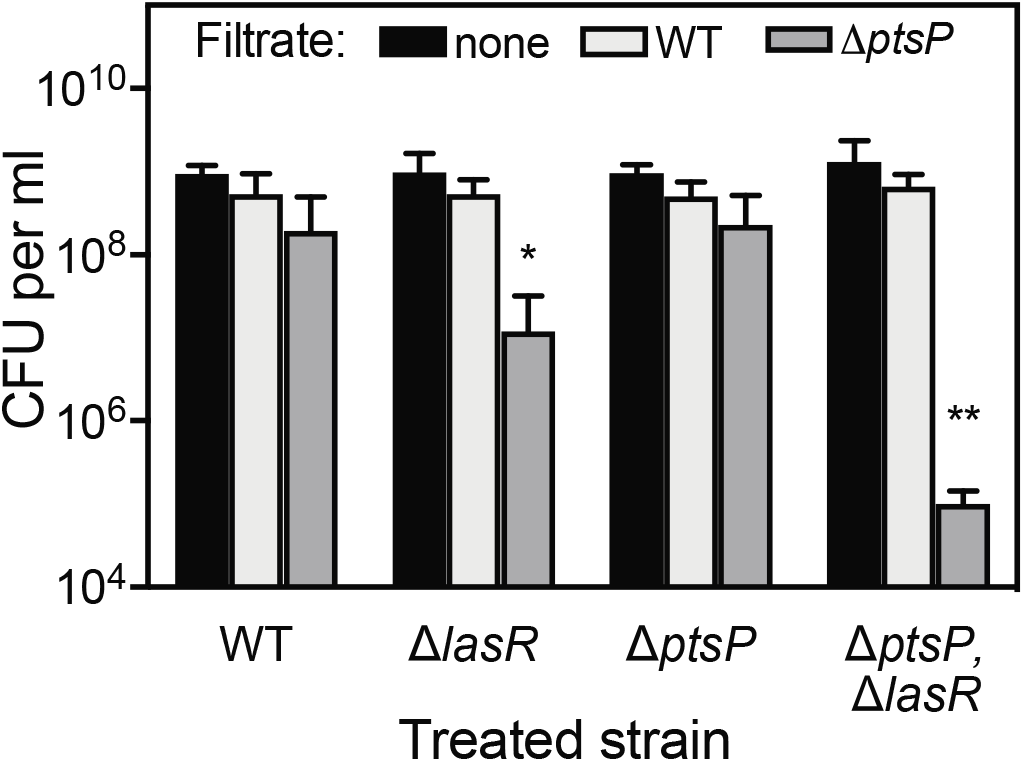
Δ*ptsP* mutant cultures has antimicrobial activity against *lasR* mutants. Final cell densities of *P. aeruginosa* strains treated for 24 h with filtered fluid from 1% casein-grown cultures of untreated (black), wild-type PA14 (light grey), or the *ptsP* mutant (dark grey). The initial cell density of the treated cultures was 5–7 × 10^5^ cells ml^-1^. The values represent the average of three independent experiments and the error bars represent the standard deviation. Statistical analysis by two-way ANOVA and Dunnett’s multiple comparison test with untreated for each strain. * p<0.05, ** p<0.01.

We hypothesized that the activity in the Δ*ptsP* mutant culture fluid was due to pyocyanin, based on our result that *ptsP* mutations increase pyocyanin production (Fig. 3). Pyocyanin is produced from the *phzA1-G1* (*phz1*) and *phzA2-G2* (*phz2*) gene clusters and the *phzM* and *phzS* genes (Fig. 6A) (45, 46). Several of the pyocyanin intermediates are toxic, such as 5-methylphenazine 1-carboxylic acid (5-Me-PCA) (47–49). We tested several gene mutations in this pathway to determine whether any of the products have activity against *lasR* mutants. We deleted *phzM*, which is needed to convert the precursor phenazine-1-carboxylic acid (PCA) to 5-Me-PCA, *phzS* which is needed to convert 5-Me-PCA to pyocyanin (45, 50), and *phzH* which is not needed for pyocyanin production but involved in production of another product of this pathway, phenazine-1-carboxamide (PCN) (45).

**Fig. 6.**
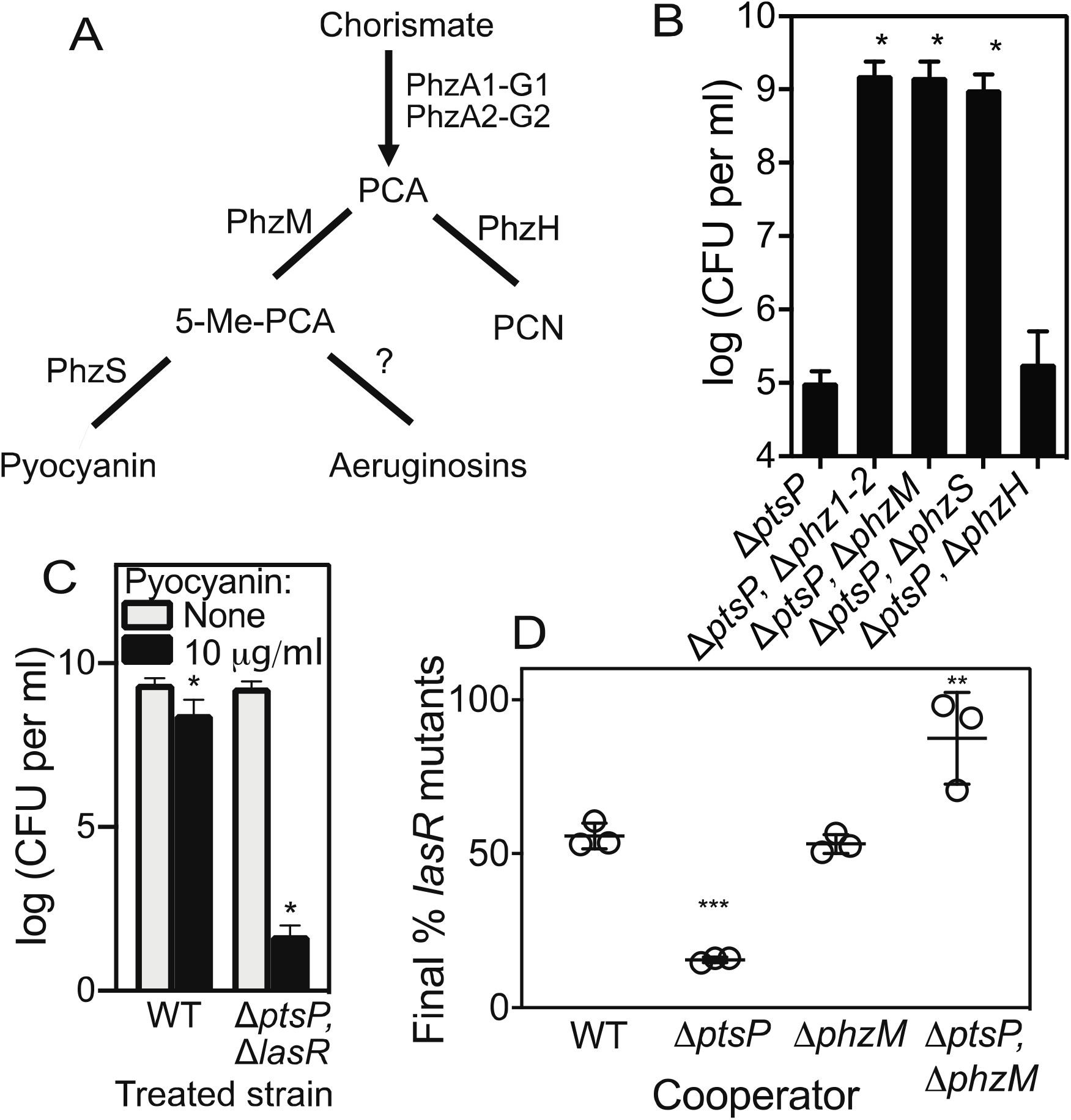
Δ*ptsP* mutants police cheaters using pyocyanin. **A.** Pyocyanin biosynthesis steps. The products of the *phzA1-G1* (*phz1*) and *phzA2-G2* (*phz2*) gene clusters synthesize phenazine-1-carboxylic acid (PCA) from chorismate. PhzM and PhzH convert PCA into 5-methylphenazine 1-carboxylic acid (5-Me-PCA) or phenazine-1-carboxamide (PCN), respectively. PhzS converts PCA into pyocyanin. **B.** Final cell densities of Δ*ptsP*, Δ*lasR* cells treated with filtered fluid from 1% casein-grown cultures of a Δ*ptsP* mutant or Δ*ptsP* with the entire *phz1* and *phz2* operons deleted (Δ*phz1-2*) or with *phzM, phzS*, or *phzH* deleted. The initial cell density of the treated cultures was 5–7 × 10^5^ cells ml^-1^. Data are the average of three independent experiments with standard deviation. Statistical analysis by one-way ANOVA and Tukey’s multiple comparisons test with Δ*ptsP*, *p<0.05. **C.** Final cell densities of wild-type PA14 (WT) or the Δ*ptsP*, Δ*lasR* mutant treated with 0 or 10 μg ml^-1^ pyocyanin and filtered fluid from 1% casein-grown Δ*ptsP* Δ*phzS* cultures. The initial cell density of treated cultures was 4–6X10^5^ CFU ml^-1^. Data are the average of four independent experiments with standard deviations. Statistical analysis by two-way ANOVA and multiple comparisons test with no-pyocyanin control, * p<0.05. **D.** Cheater suppression during competition. Competitions were inoculated with the *lasR* mutant and each cooperator strain at an initial ratio of 1:99 (cheater:cooperator) in 1% casein broth and transferred to fresh medium daily for 3 days. On day 3, cheaters were enumerated by patching as described in Materials and Methods. Each data point represents an independent experiment. The horizontal line represents the mean and the vertical line represents the standard deviation of all the experiments in each set. Statistical analysis by one-way ANOVA with Tukey’s multiple comparisons test of each set with the wild-type cooperator experiments: ** p<0.01, *** p<0.001. Total population densities of each experiment are shown in Fig. S6.

We initially tested whether pyocyanin biosynthesis contributes to the antimicrobial activity of a *ptsP* mutant by making a Δ*ptsP* Δ*phz1* Δ*phz2* triple mutant. We compared antimicrobial activity in culture fluid of this strain with that of the single Δ*ptsP* mutant, and tested against the Δ*ptsP, ΔlasR* mutant. Deleting both *phz1* and *phz2* abolished the antimicrobial activity observed with the Δ*ptsP* single mutant (Fig. 6B). Mutations in *phzM* or *phzS* similarly abolished this activity (Fig. 6B). However, no effects were observed by deleting *phzH* (Fig. 6B), which is important for production of the pyocyanin bi-product PCN but not pyocyanin itself. These results support that the antimicrobial activity in Δ*ptsP* mutant culture fluid is due to pyocyanin. We also tested whether commercially supplied pyocyanin could restore antimicrobial activity of the Δ*ptsP* Δ*phzS* mutant by adding 10 μg ml^-1^ pyocyanin to Δ*ptsP* Δ*phzS* culture filtrates prior to treating Δ*ptsP ΔlasR* cells. Adding pyocyanin reduced Δ*ptsP*, Δ*lasR* mutant growth ~10^7^-fold, whereas PA14 growth was only reduced by ~10-fold (Fig. 6C). Together, these results support the conclusion that pyocyanin can selectively limit growth of Δ*ptsP, ΔlasR* mutants.

### Pyocyanin production leads to enhanced policing by Δ*ptsP* mutants

We also tested the role of pyocyanin in policing cheaters using competition experiments. We mixed the Δ*lasR* mutant with a cooperator starting at a 1:99 (cheater:cooperator) ratio in 1% casein media and transferred the population daily for 3 days. The results are shown in Fig. 6D. In cocultures with PA14 as the cooperator, Δ*lasR* mutants increased from 1% to a final frequency of 56%. However, the Δ*lasR* mutants increased to only 16% final frequency in identical competition experiments with the Δ*ptsP* mutant, which is consistent with limited cheater proliferation observed with this strain (Fig. 4B). Deleting *phzM* in PA14 did not alter the outcome of the Δ*lasR* mutant frequency; however, when *phzM* was deleted in the Δ*ptsP* mutant cooperator, cheaters increased to a final frequency of 88%. Altogether, our results support that the cheater restriction observed in the Δ*ptsP* mutant is dependent on pyocyanin.

## DISCUSSION

This study adds to the evidence that quorum sensing contributes to antibiotic resistance in Proteobacteria. Results from these studies show that *P. aeruginosa* LasR increases tobramycin resistance in planktonic PA14 cultures. The results also show that tobramycin can limit the proliferation of *lasR* mutant cheaters in cooperating populations grown on casein. Cheaters are suppressed directly by tobramycin because they are more susceptible to growth-inhibiting effects. Cheaters are also suppressed, indirectly, by tobramycin-selected adaptive mutations that led to pyocyanin-dependent policing. Together, these results support the idea that antibiotics influence quorum sensing stability directly and indirectly through adaptive changes in the population.

These results also provide an example of how adaptive mutations can alter cheater control mechanisms. Adaptive mutations have been shown to influence cooperation and cheating in other studies (51–53). For example, adaptive mutations can improve the ability of cooperators to outcompete cheaters (19, 52) or cause re-wiring of cooperative traits and allow the cheaters to cooperate again (51). In this study, once adaptive mutations emerge in the population they caused suppression of cheaters through policing. Thus, antibiotic exposure stabilized cooperation through heritable changes that led to policing. These antibiotic-selected effects on cheating could have important implications for understanding the long-term consequences of antibiotic usage for treating infections.

In our study, cheater suppression was caused by increased production of the toxin pyocyanin. Pyocyanin was previously shown to be involved in policing *lasR* mutant cheaters (17, 54). Pyocyanin leads to the generation of highly toxic hydroxyl radicals (55–57), which can cause oxidative stress and cell death (58, 59). LasR mutant cheaters fail to upregulate enzymes involved in relieving oxidative stress such as catalase and superoxide dismutase (SOD) (17, 56, 58, 60), which explains why these mutants might be more sensitive to pyocyanin than the wild type (60, 61). However, these detoxifying enzymes can benefit both producing and non-producing members of the population (19), which may explain how *lasR* cheaters were still able to increase in frequency when cocultured with the cooperator, despite the policing effect of pyocyanin (Fig. 6). It will be interesting to determine whether pyocyanin production could have similar effects on stabilizing quorum sensing in natural communities, such as infections.

Pyocyanin production was increased due to mutations in *ptsP*. PtsP, along with two other enzymes PtsN and PtsO, make up a poorly understood system called the nitrogen phosphotransferase system (PTS^Ntr^). In *Escherichia coli*, PTS^Ntr^ is involved in regulating diverse physiological changes in response to nitrogen starvation (62, 63). It is unclear if PTS^Ntr^ plays a similar role in *P. aeruginosa*. In *P. aeruginosa*, PtsP contributes to pathogenesis (64), biofilm formation (27), and quorum sensing regulation (28). Mutations in *ptsP* have also been shown to contribute to tobramycin resistance (65, 66), consistent with our results. The specific effects of PtsP on pathogenesis, biofilm formation, and tobramycin resistance are as-yet unknown. In the case of quorum sensing, it is thought that PtsP somehow represses LasR through the antiactivator QscR (28), although it is not clear if PtsP acts entirely through QscR or modifies quorum sensing through other pathways. The important effects of mutating PtsP on *P. aeruginosa* virulence and virulence-associated behaviors suggest PtsP and the PTS^Ntr^ system might have potential as a new target for therapeutic development.

Our results also show that quorum sensing has a small but appreciable contribution to antibiotic resistance in *P. aeruginosa* strain PA14 under planktonic conditions (Fig. 1). Similar results have been reported for other strains and species, for example in *P. aeruginosa* PAO1 in biofilms (20–23), and *C. subtsugae* (10). In *C. subtsugae*, resistance is attributed to an efflux pump, CdeAB-OprA, which is transcriptionally activated by the CviR quorum-sensing receptor in response to the cognate quorum-sensing signal *N*-hexanoyl-homoserine lactone (10, 41). In *P. aeruginosa*, there may be multiple factors contributing to antibiotic resistance. There are at least three efflux pumps known to have overlapping specificity for aminoglycosides; MexAB (67, 68), MexXY (69, 70), and PA1874-1877 (71); there are also aminoglycoside-inactivating enzymes (67). Quorum-control of antibiotic resistance may provide an important evolutionary benefit. For example, quorum sensing may increase resistance to protect against self-produced toxins, or synchronize resistance factor expression across members of the population (72) to protect neighboring cells from exported antibiotic (73). Understanding how and why quorum sensing contributes to antibiotic resistance will provide important new information about the biology of quorum sensing and will be relevant to designing new therapies that function by blocking quorum-sensing systems.

## ACKNOWLEDGEMENTS

This work was supported by the NIH through grant R35GM133572 to JRC and grant GM125714 to AAD, a pilot project from the Chemical Biology of Infectious Diseases (P20 GM113117) program to JRC, and a KU Inez Jay award to JRC. Sequencing core facility support was provided by the NIH COBRE Center for Molecular Analysis of Disease Pathways Program (P20 GM103638). JHK was supported by an NIH postdoctoral fellowship (T32 AI007343). VDC was supported by an Undergraduate Research Award from the KU Center for Undergraduate Research and a K-INBRE fellowship (P20 GM103418). RGA was supported by the Fulbright Foreign Student Program (15160174). The authors would also like to acknowledge Matthew Parsek and Dingding An (University of Washington), Matthew Cabeen (Oklahoma State University) and Lars Dietrich (Columbia University) for providing *P. aeruginosa* strains and plasmids, as well as Tony Ma, Matthew Johnson, and Bryan Murphy for their technical support.

## AUTHOR CONTRIBUTIONS

R.G.A. contributed to the conception and design of the work, the acquisition, analysis and interpretation of data, and drafting and revising the manuscript. J.H.K. contributed to the design of the work, the acquisition, analysis and interpretation of data and revising the manuscript. B.M.M. contributed to the acquisition and analysis of data. V.D.C. contributed to the acquisition and analysis of data. N.E.S. contributed to the analysis and interpretation of data. A.A.D. contributed to the analysis and interpretation of data. J.R.C. contributed to the design of the work, analysis and interpretation of data, and drafting and revising the manuscript.

