## Supplemental file for "Tobramycin adaptation enhances policing of social cheaters in *Pseudomonas aeruginosa*" (from another version of this preprint: version 1, 2020-11-15)

**Supplementary Information for Abisado, et al.**

**Table S1. Bacterial strains used in this study.**

Strain	Relevant properties	Reference or source

*P. aeruginosa*

UCBPP-PA14	Ancestral wild type	(1)

PA14 Δ*lasR*	PA14 with a deletion of *lasR*	This study

PA14 Δ*rhlR*	PA14 with a deletion of *rhlR*	This study

PA14 Δ*lasR* Δ*rhlR*	PA14 Δ*lasR* with a deletion of *rhlR*	This study

PA14 Δ*lasI*	PA14 with a deletion of *lasI*	This study

PA14 Δ*ptsP*	PA14 with a deletion of *ptsP*	This study

PA14 Δ*ptsP* Δ*lasR*	PA14 Δ*ptsP* with a deletion of *lasR*	This study

PA14 Δ*phzM*	PA14 with a deletion of *phzM*	(2)

PA14 Δ*ptsP* Δ*phzM*	PA14 Δ*ptsP* with a deletion of *phzM*	This study

PA14 Δ*ptsP* Δ*phzS*	PA14 Δ*ptsP* with a deletion of *phzS*	This study

PA14 Δ*ptsP* Δ*phz1* Δ*phz2*	PA14 Δ*ptsP* with a deletion of *phz1* and *phz2*	This study

PA14 Δ*ptsP* Δ*phzH*	PA14 Δ*ptsP* with a deletion of *phzH*	This study

PA14 *fusA1* G1634A	PA14 with the *fusA1* G1634A mutation	This study

PA14 Δ*ptsP fusA1* G1634A	PA14 Δ*ptsP* with the *fusA1* G1634A mutation	This study

PA14 Δ*ptsP* CTX-1-*ptsP*	PA14 Δ*ptsP* with *attB*::CTX-1-*ptsP* insertion	This study

PA14 Δ*ptsP* CTX-1	PA14 Δ*ptsP* with *attB*::CTX-1 insertion	This study

PA14 CTX-1	PA14 with *attB*::CTX-1 insertion	This study

*P. aeruginosa* PA14 variants

*Variants isolated after daily transfer for 32 days in 1% casein*

T1	Isolate from experiment with tobramycin added at 0.6-7.1µg ml^-1^	This study

T2	Isolate from experiment with tobramycin added at 0.6-7.1µg ml^-1^	This study

T3	Isolate from experiment with tobramycin added at 0.6-7.1µg ml^-1^	This study

T4	Isolate from experiment with tobramycin added at 0.6 µg ml^-1^	This study

T5	Isolate from experiment with tobramycin added at 0.6 µg ml^-1^	This study

T6	Isolate from experiment with tobramycin added at 0.6 µg ml^-1^	This study

N1	Isolate from experiment with no added tobramycin	This study

N2	Isolate from experiment with no added tobramycin	This study

N3	Isolate from experiment with no added tobramycin	This study

*Modified variants*

T1 CTX-1	T1 Δ*ptsP* with *attB*::CTX-1 insertion	This study

T1 CTX-1-*ptsP*	T1 Δ*ptsP* with *attB*::CTX-1-*ptsP* insertion	This study

T2 CTX-1	T2 Δ*ptsP* with *attB*::CTX-1 insertion	This study

T2 CTX-1-*ptsP*	T2 Δ*ptsP* with *attB*::CTX-1-*ptsP* insertion	This study

T3 CTX-1	T3 Δ*ptsP* with *attB*::CTX-1 insertion	This study

T3 CTX-1-*ptsP*	T3 Δ*ptsP* with *attB*::CTX-1-*ptsP* insertion	This study

*Escherichia coli*

DH5α	F- Φ80*lacZ* Δ*M15* Δ(*lacZYA*-*argF*) *U169 hsdR17*(rK- mK^+^)	Invitrogen

*recA1 endA1 phoA supE44 thi-1 gyrA96 relA1* λ**^-^**

S17-1	*recA pro hsdR RP4-2-Tc::Mu-km::Tn7*	(3)

SM10	*thi thr leu tonA lacY supE recA*::RP4-2-Tc::Mu Km λ*pi*r	(3)

Rho3	*thi-1 thr-1 leuB26 tonA21 lacY1 supE44 recA*,	(4)

integrated RP4-2 Tcr::Mu (λ*pir* ^+^)Δ*asd*::FRT Δ*aphA*::FRT

**Table S2. Plasmids used in this study.**

Plasmid	Relevant properties	Reference or source

pEX18Ap	Suicide vector, Ap^r^	(5)

pDA8	pEX18Ap containing *lasR* flanking sequences of *lasR,* Ap^r^	An and Parsek,	unpublished

pDA9	pEX18Ap containing flanking sequences of *rhlR*, Ap^r^	An and Parsek,	unpublished

pEXG2	Suicide vector, Gm^r^	(6)

pEXG2-Δ*lasI*	pEXG2 containing Δ*lasI* with flanking sequences, Gm^r^	(7)

pEXG2-*ptsP*	pEXG2 containing Δ*ptsP* with flanking sequences, Gm^r^	(8)

pEXG2-*fusA1* G1634A	pEXG2 containing G1634A mutation in *fusA1,* Gm^r^	This study

pPROBE-GT	Broad-host-range pVS1/p15a GFP reporter, Gm^r^	`	(9)

pBS351	pPROBE-GT with -1 through-501 5’ region of *lasI*, Gm^r^	(10)

CTX-1	mini-CTX-1, integrative tetR plasmid for *P. aeruginosa*, Tet^r^	(11)

CTX-1-*ptsP*	CTX-1 containing the *ptsP* gene, Tet^r^	(8)

**Table S3. Primers used in this study.**

Primer	Sequence

lasR-A	GCGCGC**GAATTC**AACATGGTCACCTCCAGCA

lasR-B	GCTGAGAGGCAAGATCAGAG**CTGCAG**CATAGCGCTACGTTCTTCTT

lasR-C	GCAAGAAGAACGTAGCGCTATG **CTGCAG**CTCTGATCTTGCCTCTCA

lasR-D	GCGCGC**AAGCTT**GTTACCGTCACCAGCGTCT

rhlR-A	GCGCGC**GAATTC**GCTGTTCGACGGCAGTAT

rhlR-B	GCGCGTCGAACTTCTTCTGGAT**CTGCAG**CATTGCAGTAAGCCCTGA

rhlR-C	GCTCAGGGCTTACTGCAATG**CTGCAG**ATCCAGAAGAAGTTCGACGC

rhlR-D	GCGCGC**AAGCTT**CGGACCGCAGAGAGACTA

rgaoligo74	TAATAA**AAGCTT**CTCGGTGAAAGGCAAGAAAGA

rgaoligo75	TAATAA**TCTAGA**GCTTCGGCGTATTTGGAGAA

^a^Bolded text indicates restriction sequences

^b^rgaoligo74 and 75 were used to incorporate the *fusA1* G1634A mutation. Other oligos were used for *lasR* or *rhlR* deletions as indicated by their name.

| Isolate^a^ | *lasR* mutation(s)^b^ |
| --- | --- |
| NT1D1 | 425T, C692T |
| NT1D2 | C692T |
| NT1D3 | G72A, G99A, G144A, T215G |
| NT1D4 | C692T |
| NT1D5 | C649T |
| NT1D6 | 521C |
| NT1D7 | C692T |
| NT1D8 | G142A, G193C, G287A, C692T |
| NT2D1 | G676A |
| NT2D2 | G676A |
| NT2D3 | G676A |
| NT2D4 | G676A |
| NT2D5 | G676A |
| NT2D6 | C350T |
| NT2D7 | G217T |
| NT2D8 | G676A |

**Table S4. *lasR* mutants isolated from casein evolution experiments.**

^a^Isolates were from cultures passaged with no antibiotic.

^b^Gene mutations are given by nucleotide location.

**Table S5. Mutations in variants from populations passaged with 0.6 µg ml^-1^ tobramycin.**

| Variant | Gene^a^ | | | |
| --- | --- | --- | --- | --- |
|  | *fusA1* | P*_ygdP-ptsP_^b^* | *mdpA* | others |
| T4 | A1655G | **G392228A** | ND^c^ | *psdR* |
| T5 | G1634A | **G392228A** | 979CCTG | ND |
| T6 | ND | **A392231G** | Δ316-327 | *pstS*, *ftsH*, *nuoM*,  Δ**4064707-4092544** |

^a^Bolding indicates promoter mutations and large chromosomal deletions that are given by genomic location; all other gene mutations are given by nucleotide location.

^b^Indicates DNA sequence in the predicted promoter of *ygdP,* which is immediately upstream of *ptsP*. *ygdP* and *ptsP* are predicted to be co-transcribed using the DOOR database for prokaryotic operons (12), accessed through the website pseudomonas.com (13).

^c^ND: no mutations detected

| Strain | Doubling Time (min^-1^) | | |
| --- | --- | --- | --- |
|  | LB-MOPS | Casein broth | LB-MOPS-15 µg ml^-1^ gentamicin |
| WT | 39.35 ± 1.7 | 78.77 ± 15 |  |
| Δ*ptsP* | 39.23 ± 0.56 | 62.16 ± 8 |  |
| WT P*lasI*-*gfp* |  |  | 35.98 ± 0.57 |
| Δ*ptsP* P*lasI*-*gfp* |  |  | 35.87 ± 1.12 |
| T1 P*lasI*-*gfp* |  |  | 39.74 ± 1.94 |
| T2 P*lasI*-*gfp* |  |  | 37.54 ± 1.02 |
| T3 P*lasI*-*gfp* |  |  | 42.84 ± 3^b^ |

**Table S6. Doubling time of some strains used in this study^a^**

^a^Doubling times were calculated from hourly cell density measurements using optical density at 600 nm of cultures grown in LB-MOPS or CFU ml^-1^ of cultures grown in casein broth. The values shown are average of three independent experiments with the standard deviation.

^b^Significantly different from wild type P*lasI*-*gfp* by one-way ANOVA and Dunnett’s multiple comparisons test (p< 0.05).

**Supplementary Figures**

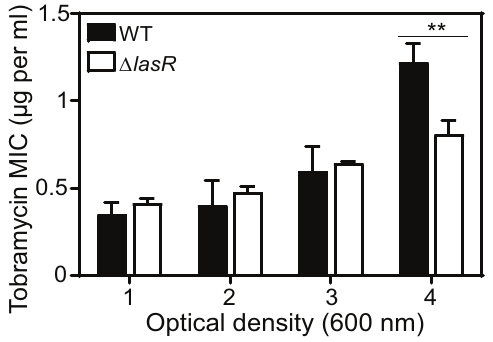

**Fig. S1.** Tobramycin resistance is LasR dependent at high cell density. The minimum inhibitory concentration (MIC) of tobramycin for *P. aeruginosa* PA14 (WT) or the PA14 Δ*lasR* mutant was determined as described in the Materials and Methods. Cultures were grown to the density indicated prior to diluting and adding tobramycin. The values represent the average of two independent experiments and the error bars represent the standard deviation. Statistical analysis by two-way ANOVA and Sidak’s multiple comparisons test with wild type: ** p<0.01.

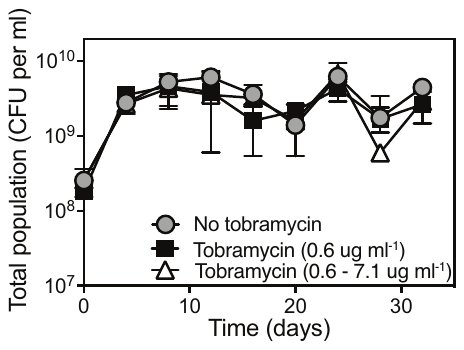

**Fig. S2.** Total cell densities *P. aeruginosa* populations passaged in casein broth. Cell densities were enumerated every 4 days by spot plating serially diluted bacteria on LB agar. *P. aeruginosa* populations were transferred daily in 1% casein broth for 32 days, and cheaters were enumerated every 4 days by patching as described in Materials and Methods. For each experiment, one culture was initially started with no tobramycin, and after 72 hr split into three cultures propagated in one of three conditions; i) continuing with no tobramycin (
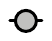
), ii) with tobramycin added at 0.6 µg ml^-1^ (
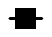
), iii) with tobramycin added at 0.6 µg ml^-1^ initially, and increasing by 50% every 4 days to a final concentration of 7.1 µg ml^-1^ (
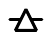
). For conditions ii and iii, tobramycin was added every other day just after transfer to fresh medium. The values represent the average of three independent experiments and the error bars represent the standard deviation.

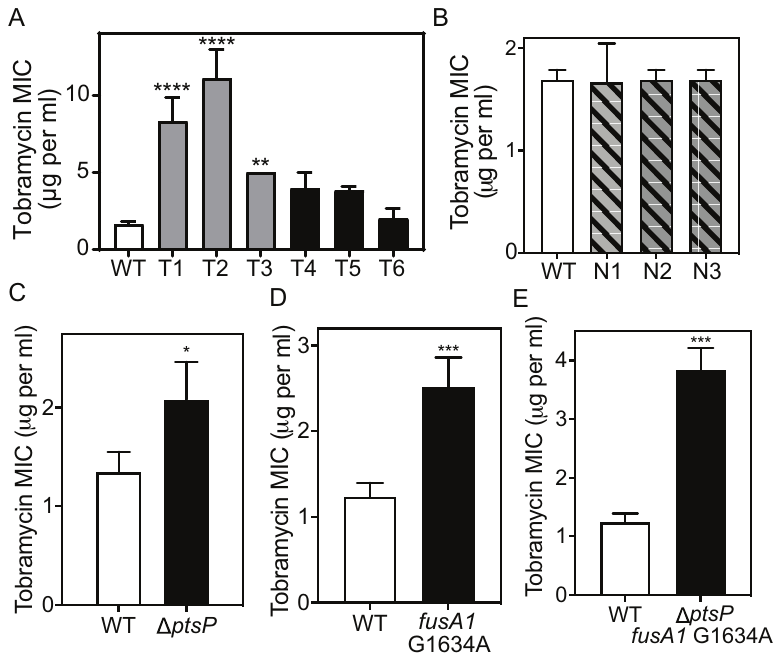

**Fig. S3.** Tobramycin susceptibility of *P. aeruginosa* mutants. **A.** The minimum inhibitory concentration (MIC) of tobramycin was determined as described in Materials and Methods. T1–3 are isolates from populations passaged in 0.6–7.1 µg ml^-1^ tobramycin while T4–6 are isolates from populations passaged in 0.6 µg ml^-1^ tobramycin. **B.** MIC of tobramycin for strains passaged in casein with no tobramycin (N1, N2, N3). **C.** MIC of tobramycin for the PA14 Δ*ptsP* mutant. **D.** MIC of tobramycin for the PA14 *fusA1* G1634A mutant. **E.** MIC of tobramycin for the PA14 Δ*ptsP*, *fusA1* G1634A mutant. The values represent the average of three (A, B, C and E) or four (D) independent experiments and the error bars represent the standard deviation. Statistical analysis by one-way ANOVA and Dunnett’s multiple comparisons test with wild type (A or B) or unpaired t-test (C, D, and E): * p<0.05, ** p<0.01, *** p<0.001, **** p<0.0001.

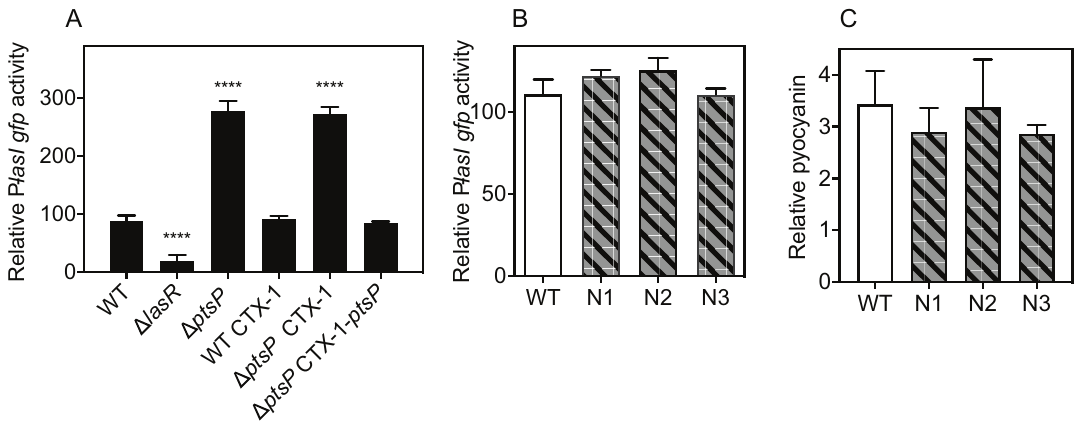

**Fig. S4.** LasR activity and pyocyanin production in *P. aeruginosa* mutants. **A** and **B.** LasR activity. *P. aeruginosa* strains were electroporated with a plasmid containing a LasR-responsive GFP reporter (pBS351). Reported values are fluorescence normalized to culture density at 18 h of growth. **C.** Pyocyanin production. Cultures were inoculated into pyocyanin-producing media, grown for 18 h and extracted before quantifying pyocyanin as described in Materials and Methods. Reported values are ug/ml pyocyanin normalized to culture density at the time of measurement. The values represent the average of three independent experiments and the error bars represent the standard deviation. Statistical analysis by one-way ANOVA and Dunnett’s multiple comparisons test with wild type, **** p<0.0001. For B and C, there was no significant difference (p>0.05).

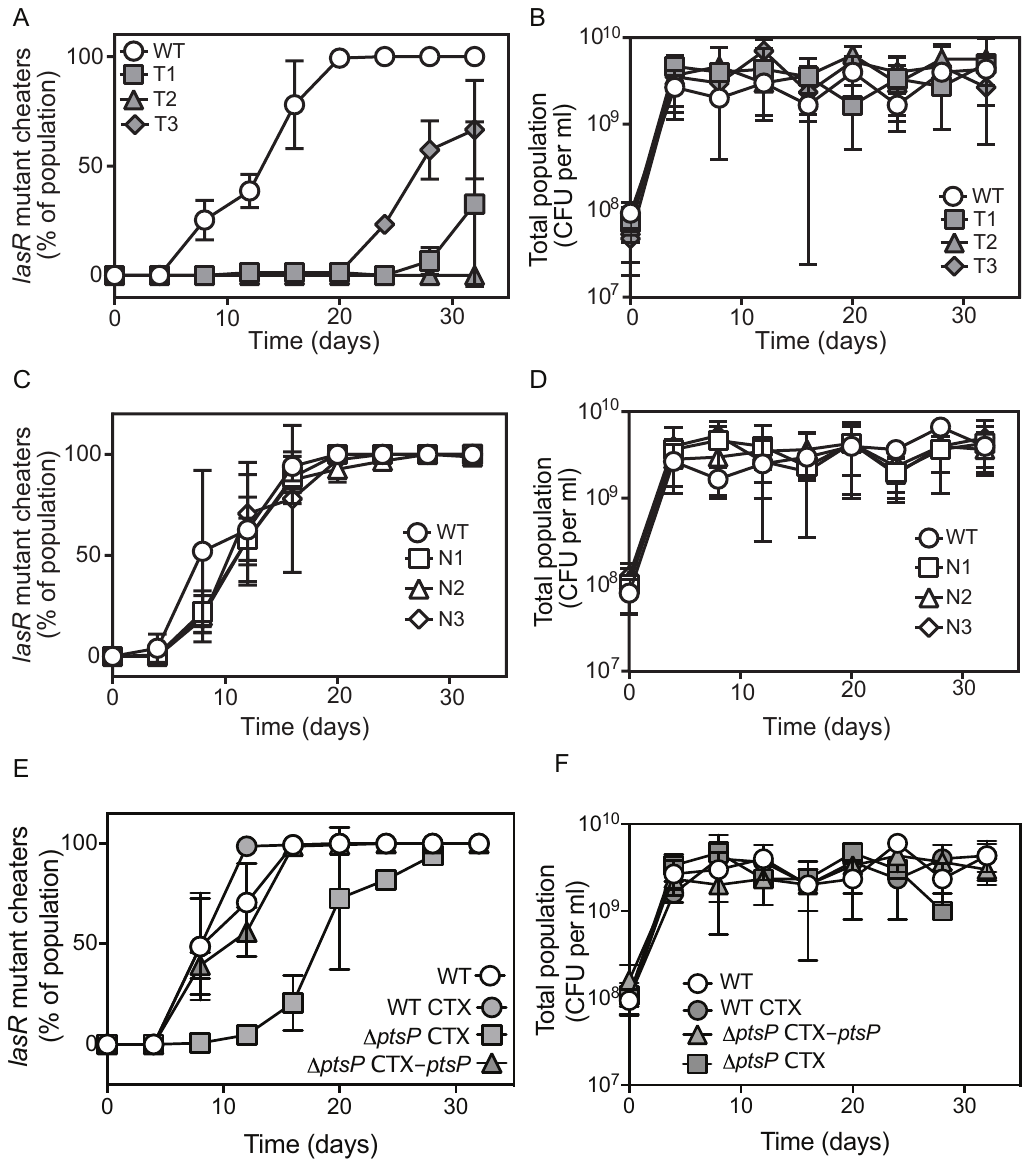

**Fig. S5.** Frequency of *lasR* mutant cheaters (A, C and E) and total populations (B, D and F) for *P. aeruginosa* strains undergoing daily passage in 1% casein for 32 days. A and B show results for tobramycin-evolved strains and corresponds with data shown in Fig. 4A of the main text. C and D show results of strains passaged with no tobramycin. E and F show results of the Δ*ptsP* strain and Δ*ptsP* carrying a complementation CTX-*ptsP* cassette and corresponds with data shown in Fig. 4B of the main text. The detection level of cheaters was 98%. The values for all graphs represent the average of three independent experiments and the error bars represent the standard deviation.

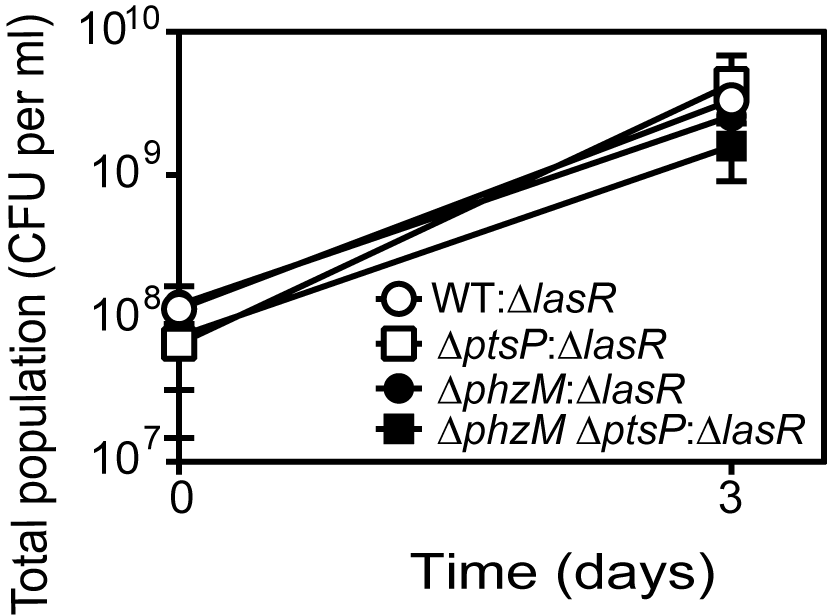

**Fig. S6.** Initial and final populations in competition experiments. Cultures were initiated at a ratio of 99:1 (cooperator:cheater) and passaged in 1% casein broth daily for 3 days. Initial and final populations were enumerated by serial dilution plating. The values represent the average of three independent experiments and the error bars represent the standard deviation. These data correspond with the data in Fig. 6D.
